# Genomic surveillance of *Neisseria gonorrhoeae* in the Philippines, 2013–2014

**DOI:** 10.1101/2020.03.19.998435

**Authors:** Manuel C. Jamoralin, Silvia Argimón, Marietta L. Lagrada, Alfred S. Villamin, Melissa L. Masim, June M. Gayeta, Karis D. Boehme, Agnettah M. Olorosa, Sonia B. Sia, Charmian M. Hufano, Victoria Cohen, Lara T. Hernandez, Benjamin Jeffrey, Khalil Abudahab, John Stelling, Matthew T.G. Holden, David M. Aanensen, Celia C. Carlos, on behalf of the Philippines Antimicrobial Resistance Surveillance Program

## Abstract

Antimicrobial-resistant *Neisseria gonorrhoeae* is a major threat to public health and is of particular concern in the Western Pacific Region, where the incidence of gonorrhoea is high. The Antimicrobial Resistance Surveillance Program (ARSP) has been capturing information on resistant gonorrhoea since 1996, but genomic epidemiology studies on this pathogen are lacking in the Philippines.

We sequenced the whole genomes of 21 *N. gonorrhoeae* isolates collected in 2013–2014 by the ARSP. The multilocus sequence type, multiantigen sequence type, presence of determinants of antimicrobial resistance and relatedness among the isolates were all derived from the sequence data. The concordance between phenotypic and genotypic resistance was also determined.

Ten of 21 isolates were resistant to penicillin, ciprofloxacin and tetracycline, due mainly to the presence of *blaTEM* gene, the S91F mutation in the *gyrA* gene and the *tetM* gene, respectively. None of the isolates was resistant to ceftriaxone or cefixime. The concordance between phenotypic and genotypic resistance was 92.38% overall for five antibiotics in four classes. Despite the small number of isolates studied, they were genetically diverse, as shown by the sequence types, the *N. gonorrhoeae* multiantigen sequence typing types and the tree. Comparison with global genomes placed the Philippine genomes within global lineage A and led to identification of an international transmission route.

This first genomic survey of *N. gonorrhoeae* isolates collected by ARSP will be used to contextualize prospective surveillance. It highlights the importance of genomic surveillance in the Western Pacific and other endemic regions for understanding the spread of drug-resistant gonorrhoea worldwide.

## Introduction

*Neisseria gonorrhoeae* is a leading cause of sexually transmitted infections, with an estimated 78 million cases of gonorrhoea each year worldwide, including 35.2 million in the WHO Western Pacific Region (*1*). In the Philippines, the prevalence of gonorrhoea in 2002 was reported to be < 2% for both men and women (*2*), with higher rates of 7.7% and 10.8% among men who have sex with men at two different sites in 2005. (*3*)

*N. gonorrhoeae* has developed resistance to first-line antibiotics such as sulfonamides, penicillins, tetracyclines, macrolides, fluoroquinolones and early cephalosporins. Currently recommended monotherapy for gonorrhoea is limited to one last effective class of antimicrobials, the extended-spectrum cephalosporins (e.g. cefixime and ceftriaxone); however, because of the recent emergence of resistance to these drugs, dual therapy with the injectable ceftriaxone plus oral azithromycin is the recommended treatment in many countries (*4*). While resistance to azithromycin has also increased globally (*1*), resistance to the dual therapy remains low (*5*).

The increase in *N. gonorrhoeae* infections resistant to front-line antibiotics triggered a global action plan from WHO to control the spread and impact of gonococcal resistance and a call for international collaborative action, especially in the Western Pacific Region (*1*). The WHO Gonococcal Antimicrobial Surveillance Programme has operated in the Western Pacific and South-East Asian regions since 1992, but surveillance of gonococcal antimicrobial resistance (AMR) remains limited in the Asia–Pacific region (*6*). In a recent report, 18 of 21 countries in the Region reported isolates with decreased susceptibility to ceftriaxone and/or isolates resistant to azithromycin between 2011 and 2016 (*6*). The Antimicrobial Resistance Surveillance Program (ARSP) of the Philippines Department of Health has been contributing AMR surveillance data to the Western Pacific Gonococcal Antimicrobial Surveillance Programme since 1996 and did not confirm isolates with decreased susceptibility or resistance to these antibiotics during 2011–2016 (*6*), while high gonococcal resistance rates against other first-line antibiotics have long been reported (Fig. 1). Continuous surveillance is thus key to detecting potential emergence or introduction of resistance to current treatment options.

**Fig. 1.**
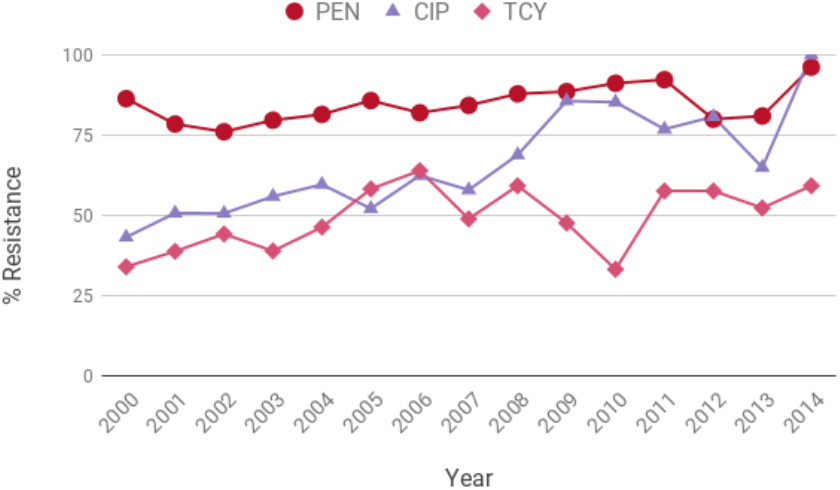
Annual resistance rates of *N. gonorrhoeae* between 2000 and 2014 for penicillin (PEN), ciprofloxacin (CIP) and tetracycline (TCY)

Molecular methods for defining the epidemiology of gonococci include both *N. gonorrhoeae* multiantigen sequence typing (NG-MAST) (*7*) and multilocus sequence typing (MLST) (*8*), although NG-MAST is more widely used to investigate specific gonococcal AMR phenotypes (*8*). Whole-genome sequencing (WGS) was recently shown to provide better resolution and accuracy than NG-MAST or MLST (*9*). Good understanding of the population structure and the mechanisms of resistance of *N. gonorrhoeae* in the Philippines would allow detection of high-risk clones associated with high-risk groups and contribute to the clinical management of gonococcal-related diseases and the creation of policies to prevent the spread of drug resistance (*10, 11*). Here, we describe the results of the first genomic survey of gonococcal isolates in the Philippines.

## Methods

### Bacterial isolates

A total of 51 *N. gonorrhoeae* isolates were collected at ARSP sentinel sites in 2013 and 2014 (Table 1). Of the 36 isolates referred to the ARSP reference laboratory for confirmation, 22 isolates from seven sentinel sites were resuscitated and submitted for WGS.

**Table 1.** Number of *N. gonorrhoeae* isolates analysed by the ARSP and referred to the reference laboratory during 2013 and 2014, isolates submitted for wholegenome sequencing and high-quality *N. gonorrhoeae* genomes obtained, by sentinel site and AMR profile.

|  | Number of Isolates |  |  |
| --- | --- | --- | --- |
|  | 2013 | 2014 | Total |
| Total ARSP | 24 | 27 | 51 |
| Referred to reference laboratory | 16 | 20 | 36 |
| Submitted for WGS | 8 | 14 | 22 |
| High-quality genomes | 8 | 13 | 21 |
| *By sentinel site*<sup>‡</sup> |  |  |  |
| CVM | 0 | 1 | 1 |
| DMC | 1 | 0 | 1 |
| EVR | 1 | 0 | 1 |
| MMH | 1 | 1 | 2 |
| NMC | 0 | 1 | 1 |
| VSM | 5 | 6 | 11 |
| ZPH | 0 | 4 | 4 |
| *By AMR profile*<sup>§</sup> |  |  |  |
| PEN CIP TCY | 2 | 8 | 10 |
| PEN CIP | 4 | 4 | 8 |
| PEN | 2 | 0 | 2 |
| CIP TCY | 0 | 1 | 1 |

‡: CVM: Cagayan Valley Medical Center, DMC: Southern Philippines Medical Center, EVR: Eastern Visayas Regional Medical Center, MMH: Corazon Locsin Montelibano Memorial Regional Hospital, NMC: Northern Mindanao Medical Center, VSM: Vicente Sotto Memorial Medical Center, ZPH: Zamboanga Del Norte Medical Center
§: PEN: penicillin, CIP: ciprofloxacin, TCY: tetracycline

### Antimicrobial susceptibility testing

All *N. gonorrhoeae* isolates in this study were tested at the ARS reference laboratory for susceptibility to five antimicrobials representing four different classes, namely penicillin (PEN), ciprofloxacin (CIP), tetracycline (TCY), ceftriaxone (CRO) and cefixime (CFX), by disc diffusion and gradient diffusion (Etest, BioMerieux). The minimum inhibitory concentrations were interpreted as resistant, intermediate or susceptible according to the interpretative criteria in the Performance Standards for Antimicrobial Susceptibility Testing (26th edition) of the Clinical and Laboratory Standards Institute (CLSI) (*12*). All isolates were screened for β-lactamase production on cefinase paper discs (BD BBL). Azithromycin was not included in the panel of antibiotics, because, at the time isolates were collected, it was not in the treatment guidelines of the Philippines and no CLSI breakpoint was available.

### DNA extraction and whole-genome sequencing

DNA was extracted from a single colony of each isolate with a Wizard^®^ Genomic DNA Purification Kit (Promega), and the quantity and quality were determined with a Quantus Fluorometer (Promega) with picogreen and a NanoDrop 2000c spectrophotometer (Thermo Fisher Scientific). The DNA extracts were then shipped to Wellcome Trust Sanger Institute for sequencing on the Illumina HiSeq platform with 100-bp paired-end reads. Raw sequence data were deposited in the European Nucleotide Archive under the study accession number PRJEB17615. Run accessions are provided on the Microreact projects.

### Bioinformatics analysis

Genome quality was evaluated according to metrics generated from assemblies, annotation files and the alignment of the isolates to the reference genome of *N. gonorrhoeae* strain TCDC-NG08107 (accession CP002441.1), as previously described (*13*). Annotated assemblies were produced with the pipeline previously described (*14*). We included 21 high-quality *N. gonorrhoeae* genomes in this study.

The MLST sequence types (STs) and NG-MAST types, as well as the presence of AMR determinants (known genes or mutations) and clustering of the isolates according to genetic similarity (tree), were predicted *in silico* from genome assemblies with Pathogenwatch (https://www.sanger.ac.uk/tool/pathogenwatch/) (*15*). In parallel, the evolutionary relations among the isolates were inferred from single nucleotide polymorphisms (SNPs) by mapping the paired-end reads to the reference genome of *N. gonorrhoeae* strain FA 1090 (accession AE004969.1), as described in detail previously (*13*). Mobile genetic elements described in the FA 1090 genome (*16*) were masked in the alignment of pseudogenomes with a script available at https://github.com/sanger-pathogens/remove_blocks_from_aln. SNPs were extracted with snp-sites (*17*), and a maximum likelihood phylogenetic tree was generated from 7518 SNP positions with RAxML (*18*), the generalized time-reversible model and the GAMMA method of correction for among-site rate variation, with 500 bootstrap replicates.

To complement the Pathogenwatch AMR results, known AMR determinants were identified from raw sequence reads with ARIBA (*19*) and a curated database of known resistance genes and mutations available at https://github.com/martinghunt/ariba-publication/tree/master/N_gonorrhoeae/Ref. The combined genotypic predictions of AMR (test) were compared with the phenotypic results (reference), and the concordance between the two methods was computed for each of five antibiotics (105 total comparisons). Isolates found to be resistant or to have reduced susceptibility (intermediate) were pooled as non-susceptible for comparison purposes. An isolate with the same outcome for both the test and reference (i.e. both susceptible or both non-susceptible) was counted as a concordant isolate. Concordance was the number of concordant isolates over the total number of isolates assessed (expressed as a percentage).

The maximum likelihood tree, genotyping results and AMR predictions and the metadata collected from the sentinel sites were visualized with Microreact (*20*).

To contextualize the genomes from this study with publicly available global genomes, we combined them with two surveys available on Pathogenwatch, a European survey of 1054 genomes (*10*) and a global survey of 395 genomes (*21*).

#### Ethics statement

Ethical approval is not applicable, as we used archived bacterial samples processed by ARSP. No identifiable data were used in this study.

## Results

### Demographic and clinical characteristics of the *N. gonorrhoeae* isolates

The 21 genomes included in this study represented seven sentinel sites, with Vicente Sotto Memorial Medical Center (VSM) contributing the most isolates (*n* = 11). The highest incidence was in the age group 15–24 years (47.6%, *n* = 10), followed by the age groups 5–14 years (28.6%, *n* = 6), 25–34 years (14.3%, *n* = 3) and 1–4 years (9.5%, *n* = 2, Table 2). The numbers of isolates from females and males were almost equal (*n* = 11 and *n* = 10, respectively). The most frequent specimen source was the vagina (*n* = 8) for female patients and penile discharge (*n* = 5) for males. All the patients were outpatients (*n* = 21).

**Table 2.** Demographic and clinical characteristics of 21 *N. gonorrhoeae* isolates.

| **Characteristic** | **No. Isolates** |
| --- | --- |
| **Sex** |  |
| Male | 10 |
| Female | 11 |
| **Age (years)** |  |
| 1–4 | 2 |
| 5–14 | 6 |
| 15–24 | 10 |
| 25–34 | 3 |
| **Patient type** |  |
| Inpatient | 0 |
| Outpatient | 21 |
| **Specimen origin** |  |
| Community | 21 |
| Hospital | 0 |
| **Specimen type** |  |
| Vagina | 8 |
| Penile discharge | 5 |
| Cervix | 3 |
| Genital discharge, male | 2 |
| Decubitus ulcer | 1 |
| Urine | 1 |
| Other | 1 |

### Concordance between phenotypic and genotypic AMR

Isolates were tested for susceptibility to five antibiotics representing four classes. In line with the resistance trends shown in Fig. 1, the most prevalent resistance profile was PEN CIP TCY, identified in 10 isolates from five sentinel sites and linked mainly to the presence of the *bla*_TEM_ gene, the S91F mutation in GyrA and the *tetM* gene, respectively (Table 3). One penicillin-resistant isolate (13ARS_DMC0024) harboured three mutations, the □57delA mutation in the *mtrR* promoter and the non-synonymous substitutions in *ponA* (L421P) and *porB* (A121D), which may contribute to high-level penicillin resistance (*22*). The partially assembled *bla*TEM gene was also detected in this genome but was not considered in the concordance analysis. We also identified the non-synonymous mutation in the *folP* gene (R228S), which confers resistance to sulfonamides, in 20 of 21 genomes; however, isolates are not routinely tested for resistance to this antibiotic class.

**Table 3.** Comparison of genomic predictions of antibiotic resistance with susceptibility testing at the ARS reference laboratory.

| Antibiotic class | Antibiotic | Resistant Isolates | False positive | False negative | Concordance (%) | Resistance genes/SNPs |
| --- | --- | --- | --- | --- | --- | --- |
| Cephalosporin | Cefixime | 0 | 0 | 0 | 100 | - |
| Cephalosporin | Ceftriaxone | 0 | 0 | 0 | 100 | - |
| Penicillin | Penicillin | 18 | 0 | 1 | 95.24 | *bla*<sub>TEM</sub>,<br>*mtrR*_promoter_-57delA, *mtrR*_A39T, *mtrR*_disrupted, *penA*_ins346D, *ponA1*_L421P, *porB1b*_A121D, *porB1b*_G120K |
| Tetracycline | Tetracycline | 14 | 7 | 0 | 66.67 | *tetM*, *rpsJ*-V57M, *mtrR*_promoter_-57delA, *mtrR*_A39T, *mtrR*_disrupted |
| Fluoroquinolone | Ciprofloxacin | 19 | 0 | 0 | 100 | *gyrA*_D95G, *gyrA*_D95A, *gyrA*_S91F, *parC*_D86N, *parC*_S87N, *parC*_E91K |

The concordance between antimicrobial susceptibility testing results and genotypic predictions (Table 3) was > 95% for all antibiotics except tetracycline (66.67%), resulting in an overall concordance of 92.38% (Table 3). The discrepancies were attributed to seven false-positive results (isolates with a susceptible phenotype but with known AMR determinants in their genomes), all of which contained the mutation V57M in the *rpsJ* gene; two isolates also carried the *tetM* gene (*23*).

### Genotypic findings

#### In silico *genotyping*

A total of 15 different STs were identified, with only four (9364, 10316, 1582 and 8133) represented by more than one isolate. Nine genomes were assigned to eight known NG-MAST types and 12 genomes to nine novel types. Only three sentinel sites, namely Corazon Locsin Montelibano Memorial Regional Hospital (MMH), VSM and Zamboanga Del Norte Medical Center (ZPH), submitted more than one isolate, and all were represented by almost as many STs as isolates submitted. Details of the numbers and the most common STs and NG-MAST types found at each sentinel site are shown in Table 4.

**Table 4.** Distribution of isolates, STs, NG-MAST types, resistance profiles and resistance genes and mutations at the seven sentinel sites. The genetic resistance mechanisms for all the isolates from each site are listed.

| Laboratory | No. of Isolates | STs (no. of isolates) | NG-MAST types (no. of isolates) | Resistance profile<sup>±</sup> (no. of isolates) | Resistance genes and SNPs |
| --- | --- | --- | --- | --- | --- |
| CVM | 1 | 8780 | Novel | PEN CIP TCY | *bla*<sub>TEM</sub>, *penA*_ins346D, *ponA1*_L421P, *mtrR*_A39T, *gyrA*_S91F, *gyrA*_D95A, *parC*_S87N, *tetM*, *rpsJ*-V57M |
| DMC | 1 | 1903 | Novel | PEN CIP TCY | *penA*_ins346D, *ponA1*_L421P, *porB1b*_A121D, *mtrR*_promoter_-57delA, *gyrA*_D95G, *gyrA*_S91F, *parC*_S87N, *parC*_E91K, *tetM*, *rpsJ*-V57M |
| EVR | 1 | 1582 | 13 796 | PEN CIP | *bla*<sub>TEM</sub>, *penA*_ins346D, *mtrR*_A39T, *gyrA*_D95G, *gyrA*_S91F, *rpsJ*-V57M |
| MMH | 2 | 11 956, 11 208 | 1631, novel | PEN CIP | *bla*<sub>TEM</sub>, *penA*_ins346D, *mtrR*_A39T, *porB1b*_A121D, *porB1b*_G120K, *gyrA*_D95G, *gyrA*_S91F, *parC*_D86N, *rpsJ*_V57M |
| NMC | 1 | 1588 | Novel | PEN CIP TCY | *bla*<sub>TEM</sub>, *penA*_ins346D, *mtrR*_A39T, *ponA1*_L421P, *gyrA*_S91F, *gyrA*_D95A, *tetM*, *rpsJ*-V57M |
| VSM | 11 | 8133 (2), 9 364 (2), 10 316 (1), 1 587 (1), 11 431 (1), 11 963 (1), 1 582 (1), 11 956 (1), 15 234 (1) | 2187(2), 1 691 (1), 1 498 (1), 1797 (1), novel (6) | PEN CIP (4)<br>PEN CIP TCY (4)<br>CIP TCY (1)<br>PEN (2) | *bla*<sub>TEM</sub>, *penA*_ins346D, *mtrR*_disrupted, *mtrR*_A39T, *ponA1*_L421P, *porB1b*_A121D, *porB1b*_G120K, *gyrA*_D95G, *gyrA*_D95A, *gyrA*_S91F, *parC*_D86N, *parC*_S87N, *parC*_E91K, *tetM*, *rpsJ*-V57M |
| ZPH | 4 | 10 316 (2), 9364 (1), 8130 (1) | 2080 (1), 11 821 (1), novel (2) | PEN CIP TCY (3)<br>PEN CIP (1) | *bla*<sub>TEM</sub>, *mtrR*_disrupted, *mtrR*_A39T, *porB1b*_A121D, *gyrA*_D95G, *gyrA*_D95A, *gyrA*_S91F, *parC*_D86N, *parC*_S87N, *parC*_E91K, *tetM*, *rpsJ*-V57M |

<sup>±</sup>PEN: penicillin, CIP: ciprofloxacin, TCY: tetracycline

#### Population structure of *N. gonorrhoeae* in the Philippines

The diverse gonococcal population was represented by a tree with three deep-branching clades and no clear geographical signal (Fig. 2). Clades II and III were characterized by a different repertoire of AMR genes and mutations. Clade II contained mostly isolates susceptible to or with reduced susceptibility to tetracycline and with the V57M mutation in the *rpsJ* gene alone, while clade III was composed of isolates resistant to tetracycline and also containing the *tetM* gene. Similarly, while both clades contained isolates resistant to ciprofloxacin harbouring the *gyrA*_S91F mutation, clade II was characterized by the presence of *gyrA_D95G* alone or in combination with the one mutation in *parC*, while clade III was characterized by the presence of *gyrA*_D95A with one or two mutations in *parC* in all but one isolates. Clades II and III showed different geographical distributions, although both were present at the VSM and ZPH sentinel sites, which submitted the most isolates. The genome from sentinel site Southern Philippines Medical Center (DMC) with a deletion in the *mtrR* promoter (□57delA) was found on a separate branch (I) in the tree and also carried a different complement of AMR determinants, indicating that it is genetically distinct from the others (Fig. 2).

**Fig. 2.**
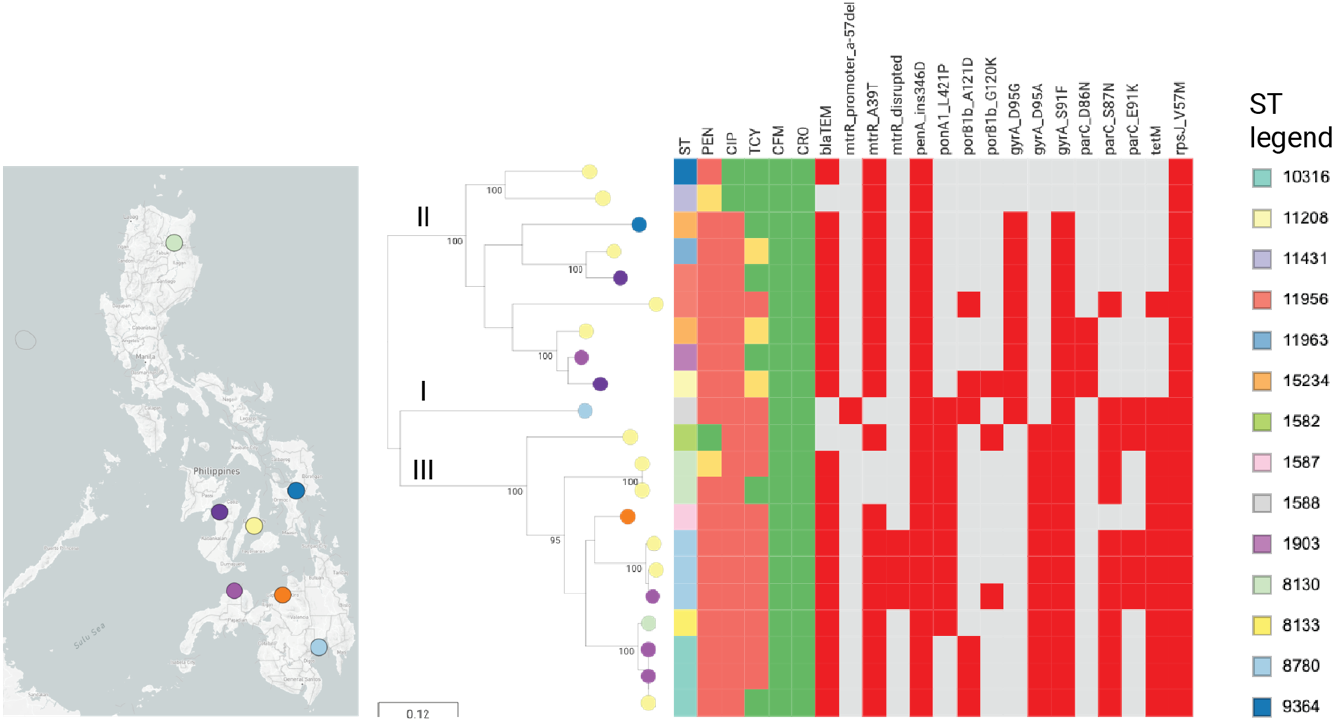
Genomic surveillance of *N. gonorrhoeae* from the Philippines 2013–2014. Maximum likelihood phylogenetic tree of 21 isolates from the Philippines, inferred from 7518 SNP sites. The tree leaves are coloured by sentinel site (map). The tree is annotated with the MLST sequence type (ST) as per the legend, the resistance phenotype for five antibiotics (red: resistant, yellow: intermediate, green: susceptible), and the distribution of known AMR mechanisms (red: present, grey: not found). The tree branches are annotated with the bootstrap values and clade designations (I, II, and III). The scale bar represents the number of SNPs per variable site. The data are available at https://microreact.org/project/ARSP_NGO_2013-14 and https://pathogen.watch/collection/flnkisnu6giu-arspngo2013-2014.

#### *N. gonorrhoeae* from the Philippines in the global context

The Philippine genomes were contextualized with two recently published collections (*10, 21*) available in Pathogenwatch. A recent global collection of 395 genomes from 58 countries (including the Philippines) showed two major lineages with different evolutionary strategies. Most of the genomes in this study were found within a subclade of lineage A, a multidrug-resistant lineage associated with infection in high-risk sexual networks (Fig. 3A), and mixed with genomes from Europe, Pakistan and South-East Asia, including genomes previously isolated in the Philippines (1998 and 2008 (*21*)). The genome from DMC was found within a separate subclade with genomes from Europe, India and Pakistan, which also carried the *mtrR*_-57delA promoter deletion.

**Fig. 3.**
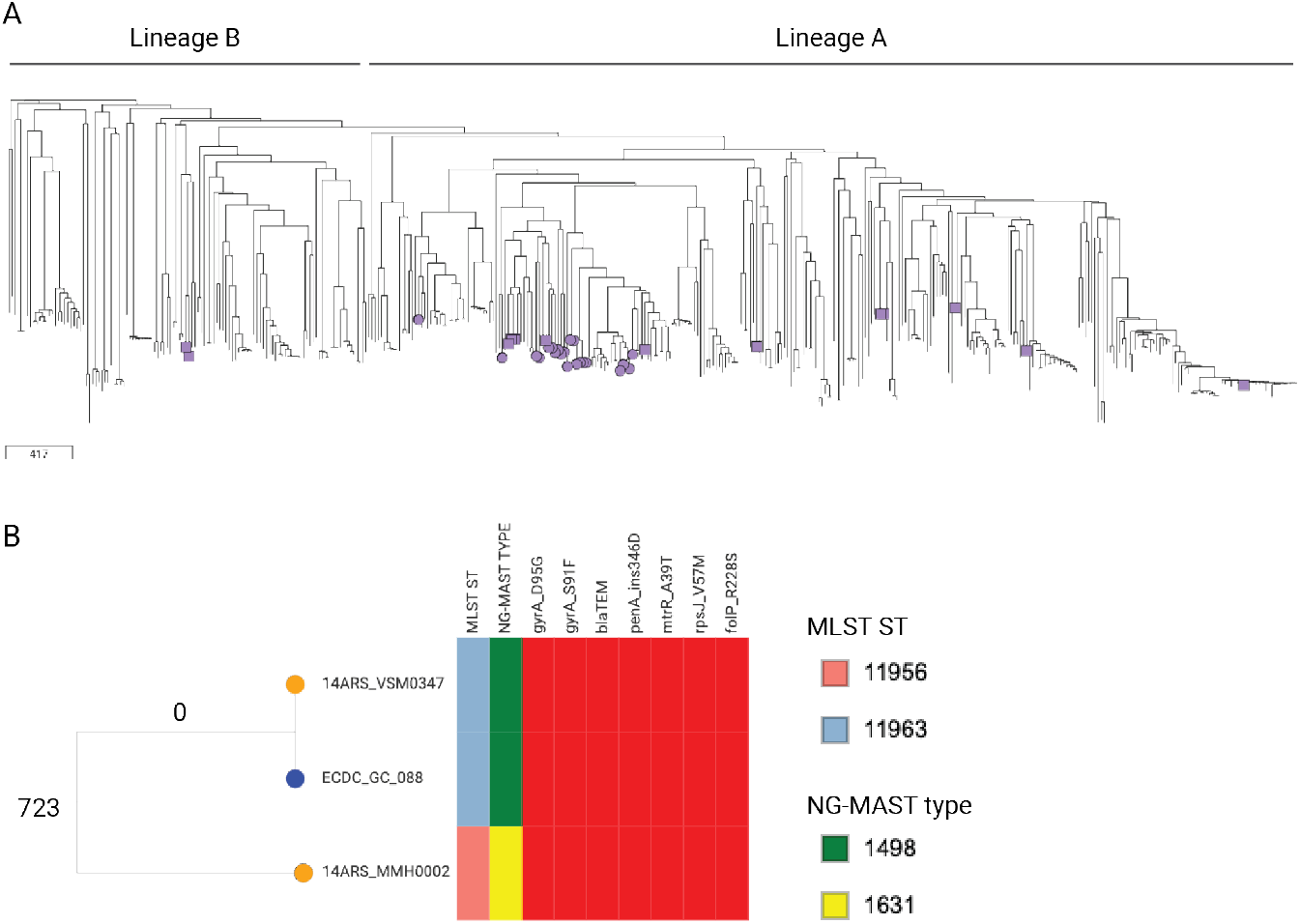
Global context of *N. gonorrhoeae* from the Philippines. **Fig. 3A.** Tree of 416 genomes from this study (*n* = 21) and a recent global collection (*n* = 395), inferred with Pathogenwatch from 22558 variant sites in 1542 core genes. Purple nodes indicate genomes from the Philippines from this study (circles) or the global collection (squares). The scale bar represents the number of SNPs. The data are available at https://pathogen.watch/collection/wrdonfwhju6f-arsp-ngo-2013-2014-global-context. **Fig. 3B.** Detail of the subtree of closely related genomes from the Philippines (orange nodes) and Norway (blue node), inferred with Pathogenwatch. The tree branches are annotated with the number of pairwise SNP differences between isolates. The metadata blocks indicate the ST, NG-MAST type and the presence (red blocks) of seven AMR determinants. The full collection is available at https://pathogen.watch/collection/xtusqgwqhxcy-arsp-ngo-2013-2014-european-context.

We further contextualized our isolates with 1054 genomes from 20 countries collected in 2013 in a European survey (*10*). Notably, the genome of strain 14ARS_VSM0347 isolated from a female on 25/04/2014 in the Philippines was highly similar to that of ECDC_GC_088 isolated from a male on 31/12/2013 in Norway (Fig. 3B), with no SNP differences found in the Pathogenwatch core genome (1542 genes). One SNP was identified in the reference-based alignment of the two pseudogenomes, confirming that these two genomes are highly similar. The two strains also shared the same complement of AMR genes and mutations (Fig. 3B). The genetic distance between the two genomes is consistent with isolates from the same location (*10*), suggesting an epidemiological link and a route of international transmission. The gonococcal reference laboratory at the Norwegian Institute of Public Health confirmed that the isolate was also resistant to penicillin and ciprofloxacin and susceptible to extended-spectrum cephalosporins (tetracycline not tested) and that the Norwegian male had visited the Philippines and claimed to have contracted gonorrhoea during his stay (personal communication).

## Discussion

WGS showed that the *N. gonorrhoeae* genomes from the Philippines are genetically diverse and carry a variety of AMR determinants, such as chromosomal mutations and acquired genes. The concordance between phenotypic and genotypic resistance was high (> 95%) for most antibiotics but only 66% for tetracycline. Susceptible isolates carrying only the *rpsJ*_V57M mutation, which confers low-level tetracycline resistance, have been reported previously (*24*). The two susceptible isolates with a full-length *tetM* gene reported by Pathogenwatch were re-tested by disc diffusion and their susceptibility confirmed. The discrepancy, which could be explained for example by the lack of expression of the gene, errors in susceptibility testing or low-level DNA contamination, will be further investigated. Although *N. gonorrhoeae* remains largely susceptible to extended-spectrum cephalosporins and azithromycin in the Philippines, we identified one isolate (13ARS_DMC0024) with a □57delA mutation in the *mtrR* promoter, which results in overexpression of the MtrCDE efflux pump and, in combination with other mutations, can increase the minimum inhibitory concentrations of azithromycin (as well as penicillin and tetracycline) (*4, 22*). Prospective surveillance with WGS can detect in real-time the acquisition of additional mutations that could result in decreased susceptibility.

A recent report suggested rapid recent intercontinental transmission of gonorrhoea, with common introductions from Asia to the rest of the world (*21*). In support of this finding, the Philippine genomes, most of which were within a subclade of global lineage A, were interspersed with those from other countries (Fig. 3A). In addition, we found evidence of an introduction event from the Philippines to Norway associated with travel of a Norwegian male to the Philippines (Fig. 3B).

Global lineage A is associated with infection in high-risk sexual networks (*21*). The combination of WGS with epidemiological information can reveal transmission routes and risk factors, which can be used to design better control measures (*25*). The small size of our retrospective data set (*n* = 21) and the linked epidemiological data did not permit any inferences about sexual networks or risk factors, which is a limitation of this study. The number of reported isolates by ARSP has, however, since increased to more than 100 per year, and data on risk factors are also collected, which will allow a more comprehensive analysis of the population diversity and of risk factors in future reports.

Our results represent the first genomic survey of *N. gonorrhoeae* isolates collected by ARSP and will constitute the background for contextualizing continuous prospective surveillance. In addition, it highlights the importance of genomic surveillance in the Western Pacific and other endemic regions to understand the spread of drug-resistant gonorrhoea worldwide.

## Acknowledgements

We are grateful to Martin Steinbakk at the Norwegian Institute of Public Health, and to Karianne Wiger Gammelsrund and the microbiology lab at the University Hospital of North Norway for retrieving the information linked to the Norwegian isolate. We also thank Dr Leonor Sánchez-Busó and Dr Magnus Unemo for critical reading of the manuscript.

